# Mechanical signaling through membrane tension induces somal translocation during neuronal migration

**DOI:** 10.1101/2023.08.28.555042

**Authors:** Takunori Minegishi, Honami Hasebe, Tomoya Aoyama, Keiji Naruse, Yasufumi Takahashi, Naoyuki Inagaki

## Abstract

Neurons migrate in a saltatory manner by repeating two distinct steps: extension of the leading process and translocation of the cell body. The former step is critical for determining the migratory route in response to extracellular guidance cues. In the latter step, neurons must generate robust forces that translocate the bulky soma against mechanical barriers of the surrounding three-dimensional environment. However, the link between the leading process extension and subsequent somal translocation remains unknown. By using scanning ion conductance microscopy, we show that leading process extension increases plasma membrane tension. The tension elevation activated mechanosensitive ion channels and triggered Ca^2+^ influx, leading to actomyosin activation at the rear of the cell. Blockade of this signaling pathway disturbed somal translocation, thereby inhibiting neuronal migration in three-dimensional environments. Thus, mechanical signaling through plasma membrane tension and mechano-channels links the leading process extension to somal translocation, allowing rapid and saltatory neuronal migration.

## INTRODUCTION

Neuronal migration is essential for brain development and neural network formation^1–5^. As a characteristic feature of neuronal migration, neurons navigate in a saltatory manner by repeating two steps^6,7^. Their migration is initiated by the extension of the leading process, which determines the migratory route in response to extracellular guidance cues^7,8^. This is followed by the translocation of the soma toward the leading process. Because the soma is the largest part of the migrating neuron, it must generate robust forces to translocate against the mechanical barriers of the surrounding three-dimensional (3D) environment^9–14^.

A previous study reported that the leading process extension of olfactory interneurons is triggered by the accumulation of shootin1b at the growth cone. This accumulation generates the traction forces necessary for the process extension^15^. On the other hand, somal translocation of cerebellar granule cells correlates with a transient increase in intracellular Ca^2+^ concentration (Ca^2+^ transient) in the soma^16^. In medial ganglionic eminence cells, Ca^2+^ transient induces actomyosin contraction at the rear of the cell, generating forces that drive somal translocation^13^. Importantly, somal translocation is initiated when the length of the leading process reaches a certain threshold^10,17^, suggesting that the leading process extension is the key step in triggering somal translocation. However, the signaling pathway that links the leading process extension and somal translocation remains unknown.

In this study, we performed live imaging of plasma membrane tension in olfactory interneurons and found that plasma membrane tension increases during the leading process extension. The tension elevation activated the mechanosensitive ion channel Tmem63b^18^ and triggered Ca^2+^ signaling, which in turn activated myosin II at the rear of the cell. Blockade of this signaling pathway by Tmem63b knockdown disturbed somal translocation and neuronal migration in 3D environments. These findings indicate that the tension-mediated signaling through mechano-channels links the extension of leading processes to the generation of robust forces for somal translocation, enabling rapid and saltatory 3D neuronal migration.

## RESULTS

### Leading process extension increases plasma membrane tension

A previous study using chick sensory neurons reported that the growth cone at the tip of neurites pulls their neurite during neurite extension^19^. In addition, recent studies reported that actin-based membrane protrusion leads to a global increase in plasma membrane tension^20–22^. To identify the signaling events triggered by leading process extension, we focused on the mechanical properties of migrating neurons. We first analyzed the membrane tension of migrating mouse olfactory interneurons in 3D Matrigel by measuring the fluorescence lifetime of the plasma membrane tension-sensitive probe Flipper-TR^23^; a longer fluorescence lifetime of the probe indicates higher membrane tension. Figure 1A shows the spatial distribution of the plasma membrane tension, which was higher in the leading process growth cone than that in the soma and leading process shaft (arrowheads, Figures 1A and 1B). Analyses of neurons bearing leading processes of different lengths demonstrated a significant positive correlation between the process length and plasma membrane tension in the soma and leading process shaft (Figures 1A and 1C).

**Figure 1.**
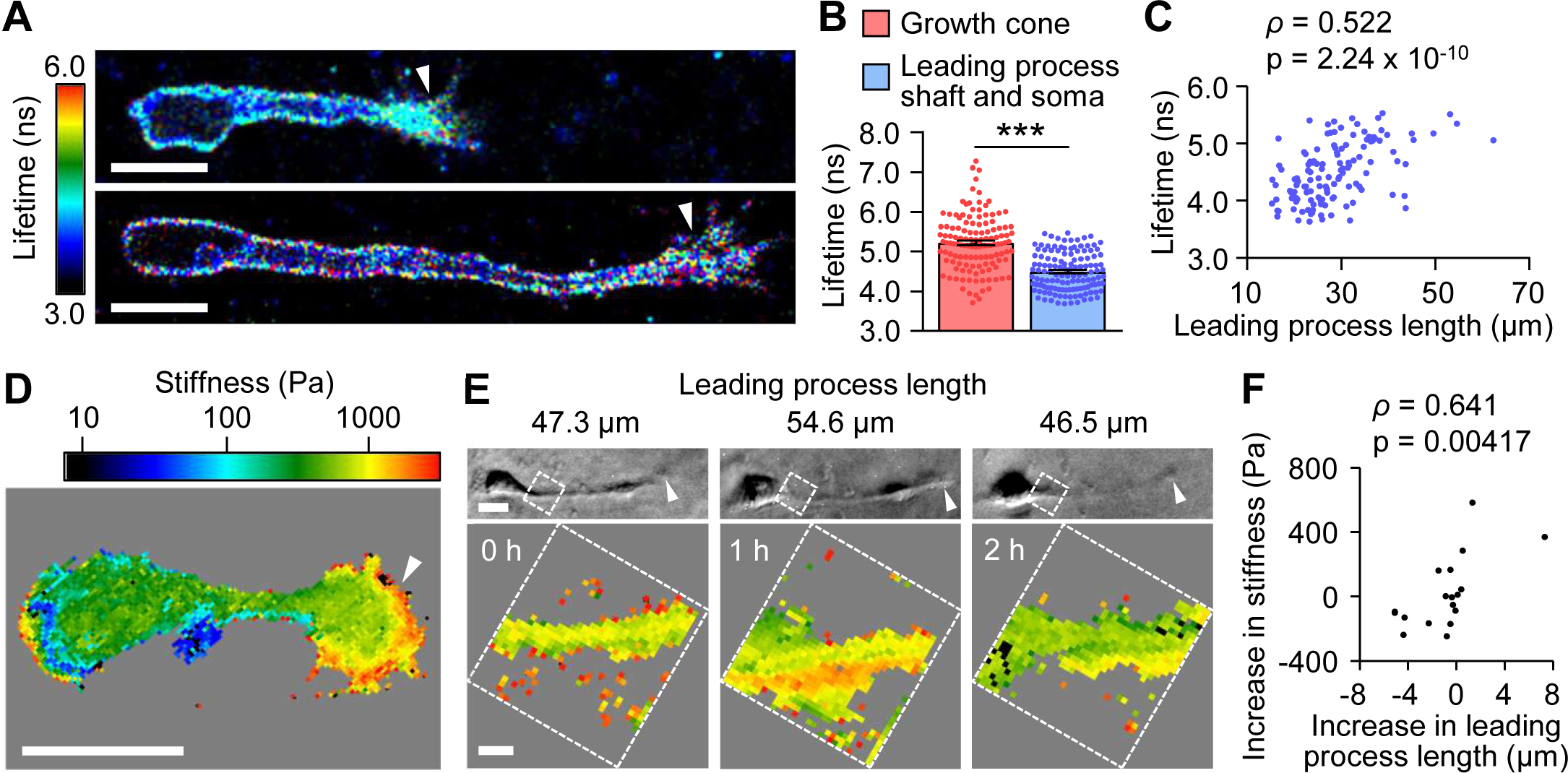
Leading process extension increases plasma membrane tension. (A) Fluorescence lifetime images of Flipper-TR in olfactory interneurons migrating in 3D Matrigel with leading processes of different lengths. The color bar indicates the lifetime in nanoseconds (ns). (B) Fluorescence lifetime of Flipper-TR measured in the growth cone or in the leading process shaft and soma. n = 129 cells. Two-tailed Wilcoxon signed-rank test. ***p < 0.01. (C) Fluorescence lifetime of Flipper-TR in the leading process shaft and soma plotted against the leading process length. n = 129 cells. Two-tailed Spearman’s correlation coefficient (*ρ*) was calculated to determine the relationship between the membrane tension and the leading process length. (D) A stiffness map of an olfactory interneuron on 2D laminin-coated substrate produced by SICM. (E) Bright field images (upper) and stiffness maps (lower) of a migrating olfactory interneuron on 2D laminin-coated substrate acquired at 1-h intervals. The arrowheads indicate the growth cone. The stiffness of the plasma membrane was measured at the proximal leading process indicated by the boxes. (F) The increase in the stiffness plotted against the increase in the leading process length. The first measurements were set as 0 as controls and omitted from the plot. n = 27 measurements from 9 cells. Two-tailed Spearman’s correlation coefficient (*ρ*) was calculated to determine the relationship between the stiffness and the leading process length. Scale bars, 10 μm (A, D and E, upper); 2 μm (E, lower).

To examine whether the leading process extension increases the plasma membrane tension, we performed time-lapse imaging. As repetitive imaging of Flipper-TR damaged neurons, we analyzed the tension with scanning ion conductance microscopy (SICM)^24^ on 2D laminin substrate. SICM has been used to monitor plasma membrane tension through cell stiffness measurements (see STAR Methods)^24,25^. Consistent with the above data, the stiffness was higher at the growth cone (arrowhead, Figure 1D) and relatively homogeneous across the leading process shaft and soma. The stiffness of the membrane in the proximal region of the leading process increased as the process extended and decreased as it shortened (Figure 1E). This relationship was confirmed by the significant positive correlation between the increased stiffness and the increase in the leading process length (Figure 1F). Overall, these data indicate that the extension of the leading process increases the plasma membrane tension.

### Increased plasma membrane tension triggers Ca^2+^ signaling through mechanosensitive ion channels

It is well established that mechanosensitive ion channels trigger cell signaling in response to extracellular mechanical stimuli^26,27^. We hypothesized that leading process extension may serve as an endogenous mechanical stimulus to trigger cell signaling. To test this possibility, we examined the relationship between leading process extension and Ca^2+^ signaling in 3D Matrigel. The intracellular Ca^2+^ concentration ([Ca^2+^]_i_) of olfactory interneurons was monitored using the Ca^2+^ indicator CalRedR525/650 (Figure 2A; Video S1). The average length of the leading process was 39.0 ± 2.7 µm (mean ± SEM, n = 14 cells), while the length of neurite extension and retraction was 11.3 ± 0.7 µm (29.0 ± 1.8% of the neurite length). Remarkably, the frequency of the Ca^2+^ transients increased when the leading process length was 4 µm larger than the average length (Figures 2B and 2C).

**Figure 2.**
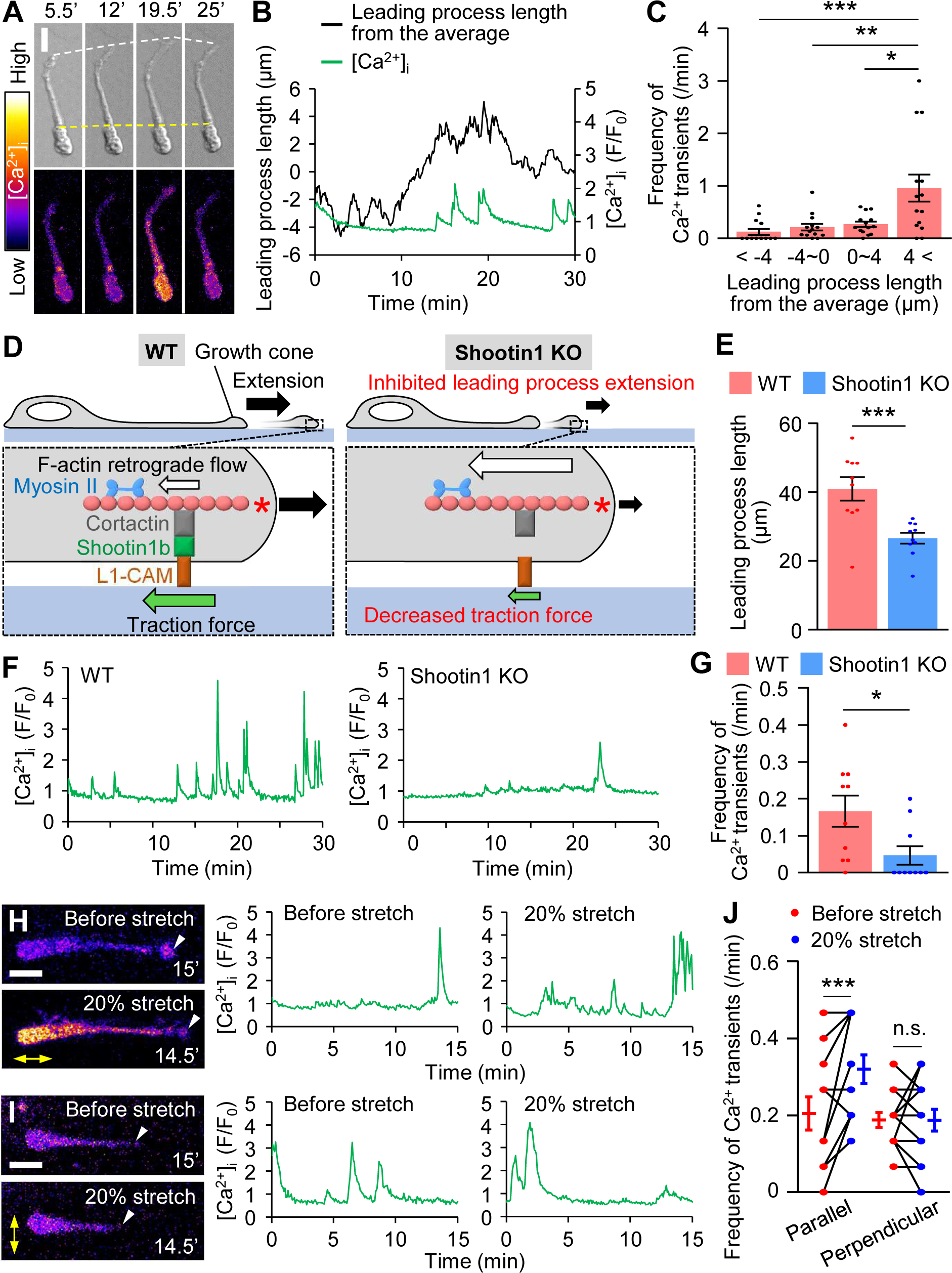
Leading process extension triggers Ca^2+^ transients. (A) Time-lapse DIC (upper) and [Ca^2+^]_i_ (lower) images of a migrating olfactory interneuron loaded with 1 μM CalRed R525/650. [Ca^2+^]_i_ is indicated as the ratio of the fluorescence intensity at 525 nm to that at 650 nm. The white and yellow dashed lines indicate the fronts of the growth cone and soma, respectively. See Video S1. (B) Time course of the leading process length and [Ca^2+^]_i_ of the neuron shown in (A). The average leading process length was set as 0. (C) Relationship between the average leading process length and the frequency of Ca^2+^ transients. The frequency of Ca^2+^ transients was measured during increases and decreases in the neurite length. n = 56 measurements from 14 cells. (D) The mechanism of shootin1b-mediated leading process extension ^15^. At the leading process growth cone, actin filaments (F-actins) undergo retrograde flow (white arrow) powered by actin polymerization (asterisk) and myosin II contraction. Shootin1b couples F-actin retrograde flow and the extracellular substrate through its interactions with an actin binding protein cortactin and a cell adhesion molecule L1-CAM. The movement of F-actin flow is transmitted through shootin1b to the substrate, thereby generating traction force on the substrate (green arrow). Traction force drives leading process extension (black arrows, left). Disruption of actin-substrate coupling by shootin1 KO decreases traction force, and inhibits leading process extension (right). (E) Leading process length of migrating WT (n = 10 cells) and shootin1 KO (n = 10 cells) olfactory interneurons. (F) Time courses of [Ca^2+^]_i_ in migrating WT (left) and shootin1 KO (right) olfactory interneurons. Neurons were imaged at 5-sec intervals for 30 min. (G) The frequency of Ca^2+^ transients in migrating WT (n = 10 cells) and shootin1 KO (n = 10 cells) neurons in (F). (H and I) Representative images of [Ca^2+^]_i_ of migrating olfactory interneurons (left) and the time courses of [Ca^2+^]_i_ (right). Neurons cultured on elastic chambers were imaged at 5- sec intervals for 15 min before (control) and after 20% stretch (yellow arrows). The arrowheads indicate the growth cones. We selected migrating olfactory interneurons with leading processes oriented parallel (H) or perpendicular (I) to the stretch direction. See Video S2. (F) The frequency of Ca^2+^ transients in migrating neurons before and after 20% stretch. Parallel, n = 11 cells; perpendicular, n = 12 cells. Data represent mean ± SEM. Statistical analysis was performed using the two-tailed Mann‒Whitney *U*-test (C), two-tailed unpaired Welch’s *t*-test (E), two-tailed unpaired Student’s *t*-test (G), two-tailed paired *t*- test (J). *p < 0.05; **p < 0.02; ***p < 0.01; n.s., no significant difference. Scale bars, 10 μm (A, H and I). See also Figure S1.

The extension of the leading process is propelled by the traction force produced by the growth cone at its tip^15,28,29^. We previously reported that shootin1b mediates the production of the traction force at the leading process growth cone^15^ (Figure 2D). Importantly, the reduction in the process length by shootin1 knockout (KO) decreased the frequency of Ca^2+^ transients (Figures 2E-2G). We also increased the leading process length using a cell-stretching device (Figure S1)^30^; increasing the process length by 20% increased the frequency of Ca^2+^ transients (Figures 2H and 2J; Video S2). On the other hand, no effect was observed when the stretch was applied perpendicular to the leading process (Figures 2I and 2J). Furthermore, treatment of neurons with 5 μM GsMTx4, which inhibits mechanosensitive ion channels^31^, reduced the frequency of Ca^2+^ transients (Figures 3A, S2A and S2B). Thus, we conclude that the increased plasma membrane tension caused by the leading process extension triggers Ca^2+^ signaling through mechanosensitive ion channels.

**Figure 3.**
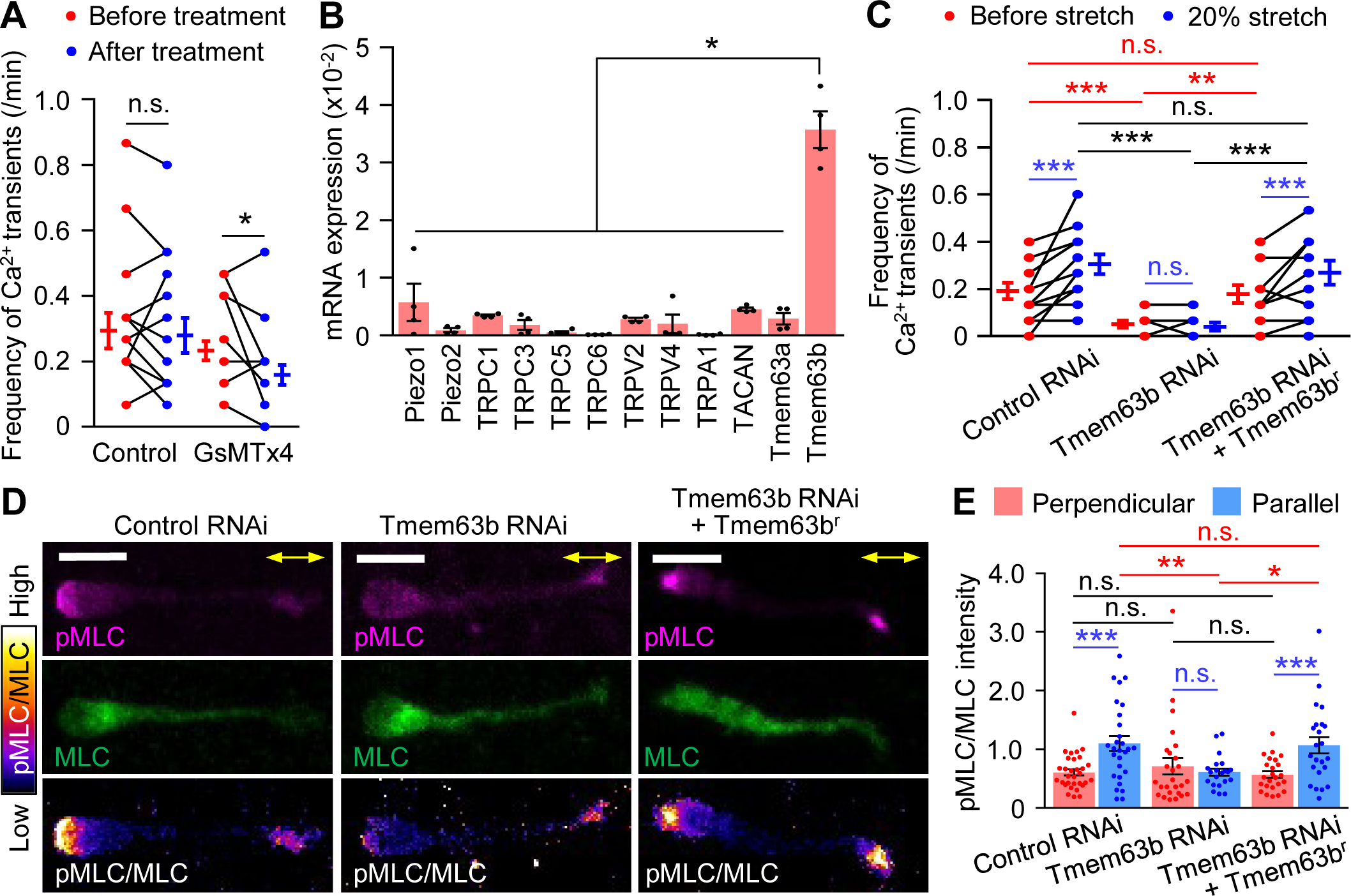
Tmem63b mediates stretch-induced Ca^2+^ transients and myosin II activation. (A) The frequency of Ca^2+^ transients in migrating olfactory interneurons before and after treatment with vehicle control (n = 12 cells) or 5 µM GsMTx4 (n = 13 cells). (B) Expression of mechanosensitive ion channels in olfactory interneurons. The mRNA of the mechanosensitive ion channels was quantified with qPCR and normalized to the expression level of GAPDH. n = 4 independent experiments. See Table S1. (C) Tmem63b knockdown abolishes the extension-triggered Ca^2+^ transient. The frequency of Ca^2+^ transients in migrating olfactory interneurons before and after 20% stretch. Control miRNA, n = 13 cells; Tmem63b miRNA, n = 12 cells; Tmem63b miRNA + RNAi- refractory Tmem63b (Tmem63b^r^), n = 12 cells. (D) Analyses of myosin II activity in olfactory interneurons expressing control miRNA, Tmem63b miRNA or Tmem63b miRNA + Tmem63b^r^. Olfactory interneurons cultured on elastic chambers were fixed after 20% stretch, and then stained with anti-phospho-myosin light chain 2 (pMLC) antibody (magenta), anti-myosin light chain 2 (MLC) antibody (green) and DAPI. The ratio of the fluorescence intensity of pMLC to that of MLC (pMLC/MLC) is displayed by the pseudocolor bar. Neurons with leading processes oriented parallel to the stretch direction (yellow arrows) were analyzed. Scale bars, 10 μm. (E) Statistical analysis of myosin II activity in olfactory interneurons from the data shown in (F). Myosin II activity (pMLC/MLC) was analyzed at the rear of the somal (Figure S3E). Control RNAi: perpendicular, n = 31 cells; parallel, n = 28 cells. Tmem63b RNAi: perpendicular, n = 26 cells; parallel, n = 21 cells. Tmem63b RNAi + Tmem63b^r^: perpendicular, n = 24 cells; parallel, n = 23 cells. Data represent mean ± SEM. Statistical analysis was performed using the two-tailed Wilcoxon signed-rank test (A), two-tailed Mann‒Whitney U-test (B and E, blue), two-tailed paired t-test (C, blue), and two-tailed one-way ANOVA with Tukey’s post hoc test (C, red and black) and two-tailed Kruskal‒ Wallis test with Dunn’s post hoc test (E, red and black). *p < 0.05; **p < 0.02; ***p < 0.01; n.s., no significant difference. See also Figures S3.

### Tmem63b mediates tension-triggered Ca^2+^ signaling

To identify the molecule involved in the tension-mediated Ca^2+^ signaling process, we analyzed the expression of mechanosensitive ion channels through quantitative PCR (qPCR) and found that Tmem63b is highly expressed in olfactory interneurons (Figure 3B). Repression of Tmem63b by miRNA suppressed spontaneous Ca^2+^ transients (Figures 3C, red and S3A-S3C) and abolished tension-triggered Ca^2+^ transients (Figure 3C, blue). Furthermore, expression of RNAi-refractory Tmem63b rescued the inhibition of spontaneous and tension-triggered Ca^2+^ transients (Figures 3C and S3D), indicating that Tmem63b is involved in tension-triggered Ca^2+^ signaling by olfactory interneurons.

### Tension-triggered Ca^2+^ signaling drives somal translocation through myosin II activation

Ca^2+^ signaling activates the myosin light chain kinase-myosin II pathway^32^. Therefore, we examined whether the activation of Tmem63b triggers actomyosin contraction and somal translocation. Immunostaining analyses of phosphorylated myosin light chain (pMLC) showed that stretching of the leading process elevated myosin II activity at the rear of the cell (Figures 3D, 3E, S3E and S3F). Furthermore, this activation was abolished by Tmem63b knockdown, indicating that tension-triggered Ca^2+^ signaling activates myosin II (Figures 3D and 3E). When Tmem63b was knocked down, neurons extended longer leading processes (Figures 4A-4C). However, despite the significant process extension, the soma did not translocate efficiently in 3D Matrigel (Figures 4A and 4B; Video S3). Neurons showed slower somal translocation speeds than control cells (Figure 4D), which was accompanied by a significant reduction in the neuronal migration speed (Figure 4E). Similar inhibition was observed in neurons treated with the MLC kinase inhibitor ML-7 (10 μM) or the myosin II inhibitor blebbistatin (100 μM) (Figure S4). Furthermore, expression of RNAi-refractory Tmem63b in neurons expressing Tmem63b miRNA rescued extension-triggered myosin II activation (Figures 3D and 3E) as well as the somal translocation and migration speed in 3D Matrigel (Figures 4A-4E; Video S3). Taken together, these data indicate that tension-triggered Ca^2+^ signaling drives somal translocation through myosin II activation.

**Figure 4.**
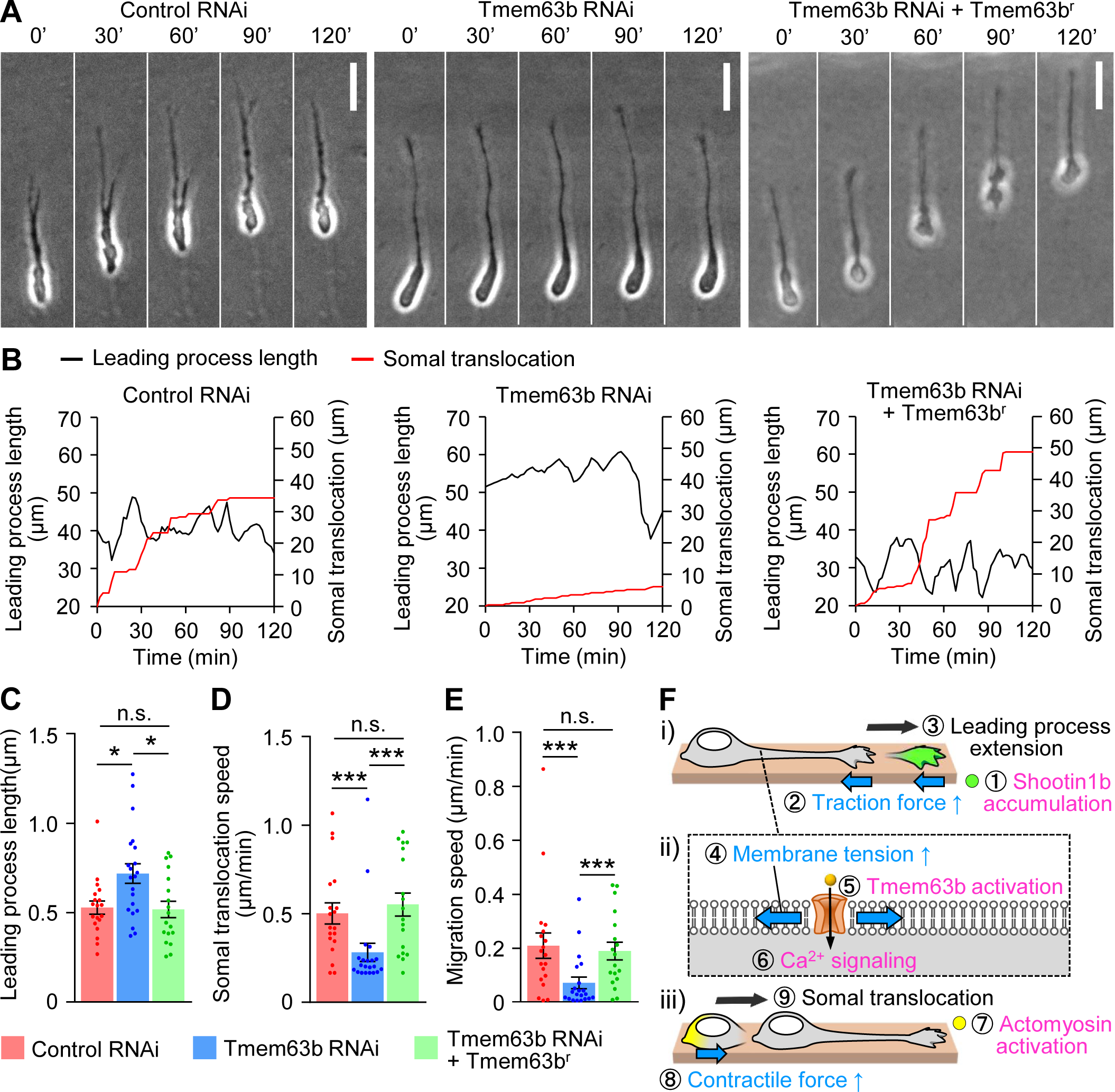
Tmem63b knockdown inhibits saltatory and efficient neuronal migration. (A) Time-lapse images of migrating olfactory interneurons expressing control miRNA, Tmem63b miRNA or Tmem63b miRNA + Tmem63b^r^. Images were acquired at 2-min intervals for 120 min. See Video S3. Scale bar, 10 μm. (B) Leading process length (black lines) and somal translocation (red lines) of olfactory interneurons shown in (A). (C - E) Leading process length (C), somal translocation speed (D) and migration speed (E) of olfactory interneurons shown in (A). Control miRNA, n = 19 cells; Tmem63b miRNA, n = 21 cells; Tmem63b miRNA + Tmem63b^r^, n = 18 cells. (F) An integrated model of saltatory neuronal migration, involving a series of biochemical (magenta) and mechanical (blue) steps (①-⑨). i) Shootin1b accumulates at the leading process growth cone (green) and couples F-actin retrograde flow and the extracellular substrate, thereby generating traction force to drive leading process extension (see Figure 2D). ii) The increase in the process length elevates the plasma membrane tension. The elevated membrane tension activates Tmem63b, which induces Ca^2+^ influx and triggers Ca^2+^ signaling. iii) Upon activation induced by Ca^2+^ signaling, actomyosin (yellow) produces contractile forces, pushing the nucleus forward and driving somal translocation. Data represent mean ± SEM. Statistical analysis was performed using the two-tailed Kruskal‒Wallis test with Dunn’s post hoc test (C - E). *p < 0.05; **p < 0.02; ***p < 0.01; n.s., no significant difference. See also Figure S4.

## DISCUSSION

In this study, we have shown that leading process extension of migrating olfactory interneurons increases plasma membrane tension. This tension elevation activates the mechanosensitive ion channel Tmem63b and triggers Ca^2+^ signaling, which drives somal translocation through actomyosin contraction. By providing a missing link to supplement previous findings, our work presents an integrated model of neuronal migration (Figure 4F) involving a series of biochemical (magenta) and mechanical (blue) steps (①-⑨). (i) Shootin1b accumulation at the growth cone generates the traction force and triggers leading process extension^15^. (ii) The leading process extension increases the plasma membrane tension, which activates Tmem63b and triggers Ca^2+^ signaling. (iii) Upon activation by Ca^2+^ signaling, actomyosin produces contractile forces, pushing the nucleus forward and driving somal translocation^13,16^. This is followed by shootin1b accumulation at the growth cone, initiating the next cycle.

It is widely accepted that mechanosensitive channels trigger cell signaling in response to mechanical stimuli from the extracellular environment, including cell compression, shear stress and substrate deformation^26,27^. Here we show that cells use a similar system in response to intracellular forces for cell migration. In keratocytes and neutrophils, it has been proposed that plasma membrane tension itself pushes the cell rear forward, thereby propelling cell migration^33,34^. However, our data have shown that blockade of the tension-mediated actomyosin activation by Tmem63b knockdown, as well as by the inhibition of MLCK or myosin II, disturbed somal translocation and inhibited neuronal migration in 3D Matrigel (Figures 4A-E and S4). These data suggest that plasma membrane tension is not sufficient to translocate the neuronal soma against the mechanical barriers of the surrounding 3D environment. In order to drive somal translocation in the 3D environment, neurons must generate robust forces through the tension-mediated signaling pathway.

We speculate that dysregulation of the signaling pathway through plasma membrane tension may occur in diseases caused by neuronal migration defects, including brain malformation, intellectual disability and epilepsy^1,5,35–37^. After migration, neurons undergo dynamic morphological changes to perform their activities, including, polarity formation, axon guidance, synapse formation, synaptic plasticity and neural network regeneration. Similar mechanical signaling, through membrane tension and mechanosensitive channels, could occur during these processes as they can also alter the plasma membrane tension locally or globally.

## STAR METHODS

Detailed methods are provided in the online version of this paper and include the following:

- KEY RESOURCES TABLE
- **RESOURCE AVAILABILITY**

`o` Lead Contact
`o` Materials Availability
`o` Data and Code Availability
- EXPERIMENTAL MODEL AND SUBJECT DETAILS

`o` Animals
`o` Culture of dissociated SVZ-derived olfactory interneurons and transfection
- METHOD DETAILS

`o` DNA constructs
`o` Analysis of plasma membrane tension by Flipper-TR
`o` Time-lapse imaging of plasma membrane tension by SICM
`o` Ca^2+^ Imaging
`o` Mechanical stretching of the leading process
`o` Analyses of neuronal migration in Matrigel
`o` qPCR analysis of migrating neurons
`o` Preparation of cell lysate and immunoblotting
`o` Immunocytochemical analysis
`o` Chemicals
- QUANTIFICATION AND STATISTICAL ANALYSIS

## ACKNOWLEDGMENTS

We thank Dr. Motohiro Nishida for valuable discussions; Mieko Ueda and Kazumi Maekawa for technical support; and Satoko Shimamura for kind encouragement. This research was supported in part by AMED under Grant Number 22gm0810011h0006 (N.I.), JSPS KAKENHI (JP19H03223, N.I.), JSPS Grants-in-Aid for Early-Career Scientists (JP19K16258 and JP23K14181, T.M.), and the Osaka Medical Research Foundation for Incurable Diseases (T.M.).

## AUTHOR CONTRIBUTION

T.M., H.H. and T.A. performed the experiments and data analysis. T.M., H.H., T.A., K.N., Y.T. and N.I. designed the experiments. T.M. and N.I. wrote the manuscript. All authors discussed the results and commented on the manuscript.

## STAR METHODS

### KEY RESOURCE TABLE

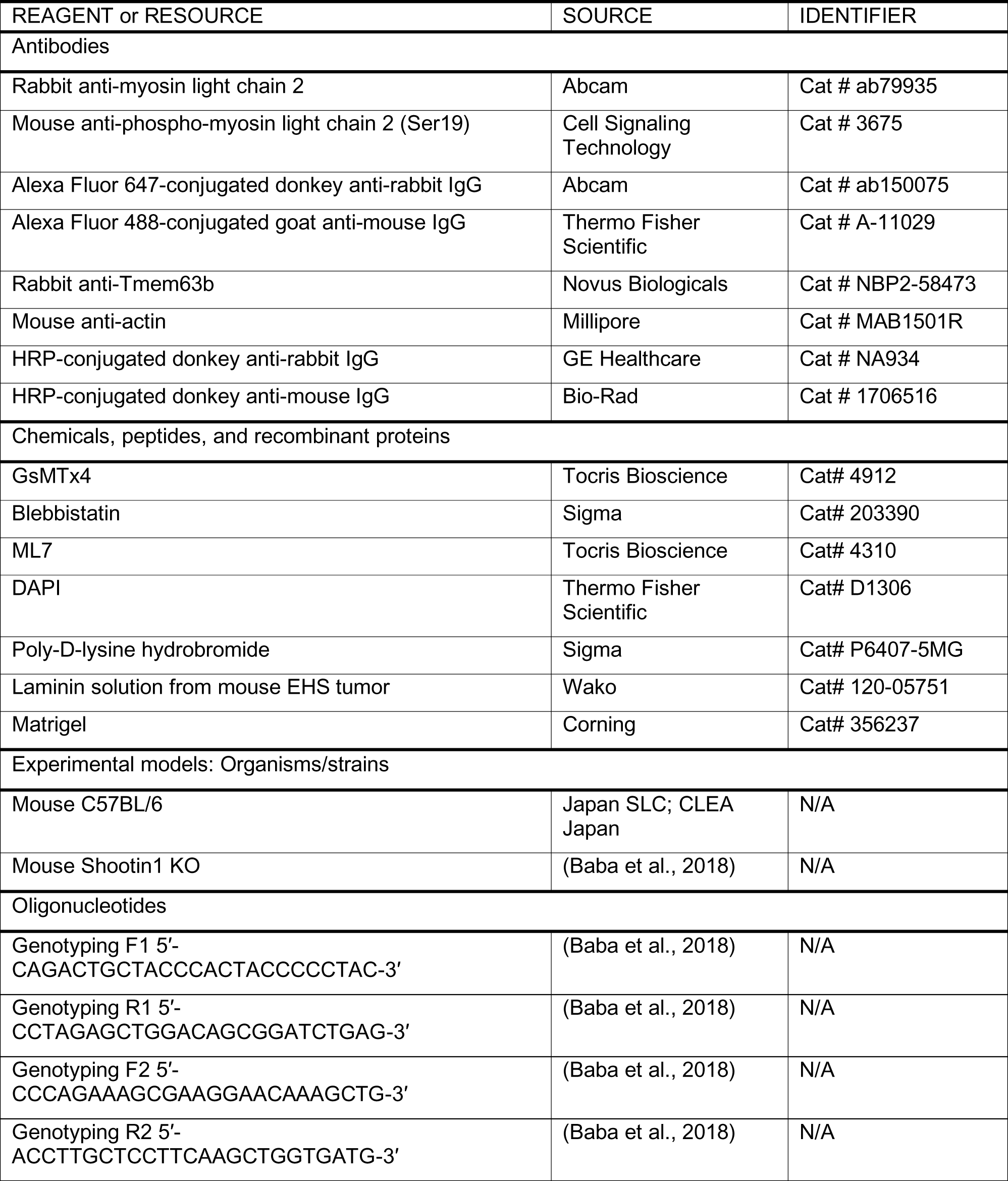
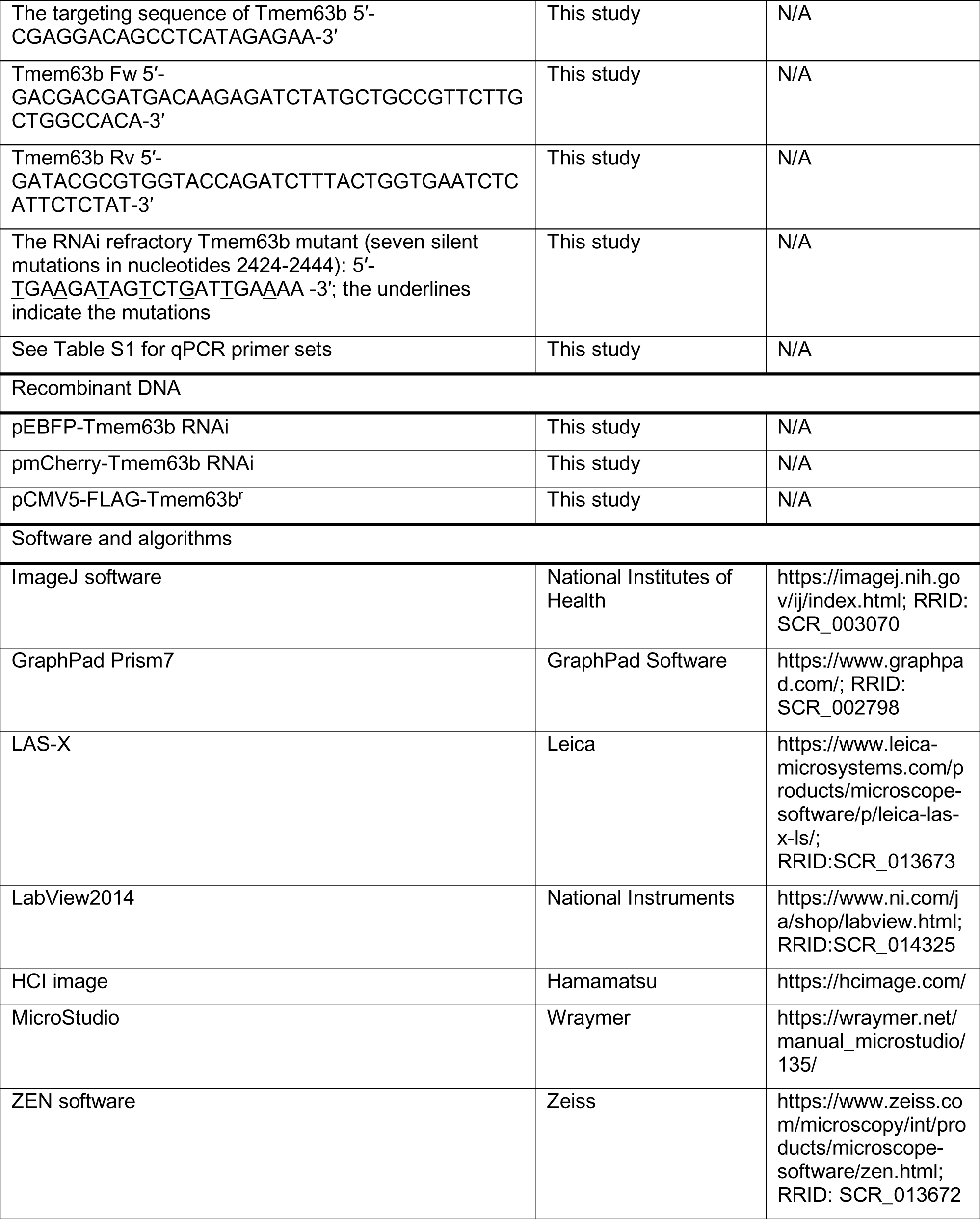
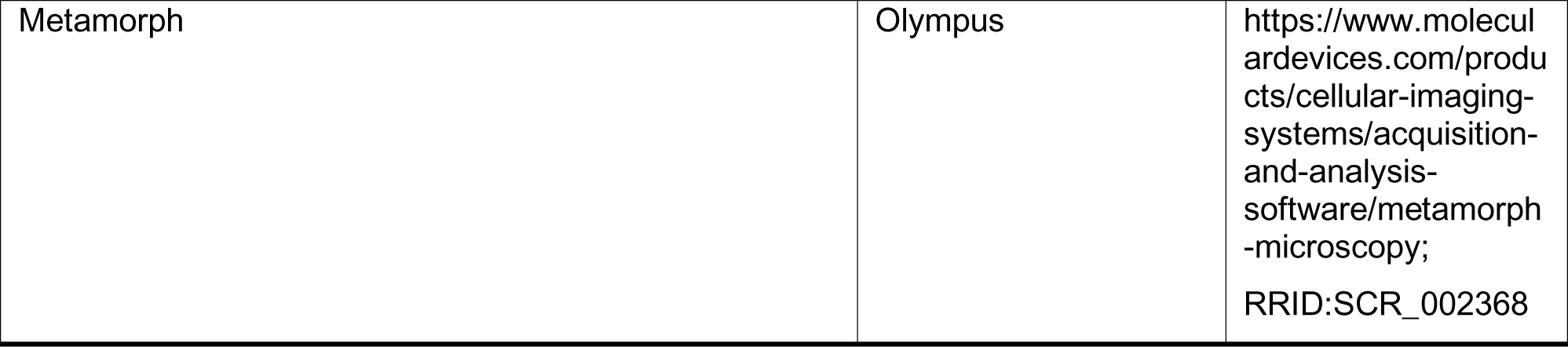

## RESOURCE AVAILABILITY

### Lead Contact

Further information and requests for resources and reagents should be directed to and will be fulfilled by the Lead Contact, Naoyuki Inagaki.

### Materials Availability

All unique materials generated in this study are available from the Lead Contact with a completed Materials Transfer Agreement.

### Data and Code Availability

- All data reported in this paper will be shared by the lead contact upon request.
- Source data of immunoblot and statistical analyses in the figures and the codes used for SICM data analyses have been deposited at Mendeley Data (10.17632/z8km7746tp.1) and are publicly available as of the date of publication.
- Any additional information required to reanalyze the data reported in this paper is available from the lead contact upon request.

## EXPERIMENTAL MODEL AND SUBJECT DETAILS

### Animals

All relevant aspects of the experimental procedures were approved by the Institutional Animal Care and Use Committee of Nara Institute of Science and Technology. Postnatal day 5 (P5) C57BL/6 mice were obtained from Japan SLC and CLEA Japan. P5 Shootin1 KO pups were obtained by crossing male and female shootin1 heterozygous mice; the offspring genotypes were checked by PCR with the following primers: Genotyping F1 (5′- CAGACTGCTACCCACTACCCCCTAC-3′) and Genotyping R1 (5′-CCTAGAGCTGGACAGCGGATCTGAG-3′) for the WT allele; Genotyping F2 (5′- CCCAGAAAGCGAAGGAACAAAGCTG-3′) and Genotyping R2 (5′-ACCTTGCTCCTTCAAGCTGGTGATG-3′) for the shootin1 KO allele. Shootin1 KO pups and their littermates were bred with their mother under standard conditions (12 h/12 h light/dark cycle, access to dry food and water). Mice of both sexes were used for experiments. The generation of shootin1 KO mice is described elsewhere ^38^. Chimeric mice were crossed with C57BL/6 mice for at least nine generations before analysis.

### Culture of dissociated SVZ-derived olfactory interneurons and transfection

SVZ tissues dissected from P5 mouse brains were dissociated with papain (Nacalai Tesque) as previously described ^15^. The dissociated olfactory interneurons were reaggregated in neurobasal medium (Thermo Fisher Scientific) containing 10% fetal bovine serum (Thermo Fisher Scientific), 2% B-27 supplement (Thermo Fisher Scientific), 1 mM glutamine (Nacalai Tesque) and 100 U/mL penicillin‒streptomycin (Nacalai Tesque) for 5-7 h. The aggregates were then embedded in a mixture of 75% Matrigel (Corning) and 25% Leibovitz′s L-15 (L-15) medium (Thermo Fisher Scientific), plated on glass bottom dishes (Matsunami) or elastic silicon chambers (Strex, catalog number STB-CH-04), and cultured in neurobasal medium containing 2% B-27 supplement, 1 mM glutamine and 100 U/mL penicillin‒streptomycin in a humidified 5% CO2 incubator at 37°C. For cell stiffness measurements by SICM, the dissociated olfactory interneurons were cultured on glass bottom dishes coated subsequently with 100 μg/mL poly-D-lysine (Sigma) and 5 μg/mL laminin (Wako Pure Chemical Industries) as previously described ^15^. For RNAi experiments, olfactory interneurons were transfected with vectors using Nucleofector (Lonza) before reaggregation. For the immunoblot analysis shown in Figure S3A, transfected olfactory interneurons were cultured on poly-D-lysine-coated plastic dishes.

## METHOD DETAILS

### DNA construction

To generate a Tmem63b miRNA expressing vector, we used a Block-iT Pol II miR RNAi expression vector kit (Thermo Fisher Scientific). The targeting sequence of Tmem63b (5′- CGAGGACAGCCTCATAGAGAA-3′, corresponding to nucleotides 2424-2444 in the coding region of mouse Tmem63b) was cloned and inserted into the pcDNA6.2- GW/EmGFP-miR expression vector. To prevent signal overlap between the GFP reporter of the miRNA expression vector and the fluorescent dyes used in our experiments, a DNA cassette containing the miRNA sequence was subcloned and inserted into pEBFP-C1 (Takara Bio) and pmCherry-C1 (Takara Bio) for Ca^2+^ imaging and immunocytochemical analyses, respectively. Full-length cDNA of mouse Tmem63b was obtained by PCR amplification of mouse subventricular zone (SVZ) cDNA with the primers Tmem63b Fw (5′-GACGACGATGACAAGAGATCTATGCTGCCGTTCTTGCTGGCCACA-3′) and Tmem63b Rv (5′- GATACGCGTGGTACCAGATCTTTACTGGTGAATCTCATTCTCTAT-3′), and subcloned and inserted into pCMV5-FLAG vector (Agilent Technology). The mouse SVZ cDNA was prepared as described in the ‘qPCR analysis of migrating neurons’ section. The RNAi refractory Tmem63b mutant was generated by inducing seven silent mutations in nucleotides 2424-2444 of Tmem63b (5′-<u>T</u>GA<u>A</u>GA<u>T</u>AG<u>T</u>CT<u>G</u>AT<u>T</u>GA<u>A</u>AA-3′; the underlines indicate the mutations) by PCR.

### Analysis of plasma membrane tension by Flipper-TR

After 2 days of culture in Matrigel, olfactory interneurons were incubated with 1 µM of the membrane tension-sensitive probe Flipper-TR (Spirochrome) diluted in culture medium for 1 h at 37°C and 5% CO_2_. Then the culture medium was replaced with L-15 medium containing 2% B-27 supplement, 1 mM glutamine and 100 U/mL penicillin‒streptomycin. Fluorescence lifetime imaging microscopy (FLIM) analysis was performed at 37°C using a confocal laser scanning microscope (SP8, Leica) equipped with a Fast Lifetime Contrast (FALCON) module, HC PL APO CS2 100x/1.40 oil immersion lens, and imaging software Las-X (Leica). The cells were excited with a pulsed 488 nm laser operating at 80 MHz, and the emission signals were detected at 575-625 nm. Following imaging, a double exponential fitting was performed to ensure a close fit (chi squared < 1.5). For quantification, we manually identified a plasma membrane region and measured the average lifetime within this region.

### Time-lapse imaging of plasma membrane tension by SICM

A membrane tension is isotropic and will resist deformation of the membrane in all directions. When plasma membrane is indented, the tension will produce a restoring force. As an increase in the plasma membrane tension leads to decreased membrane deformability ^39^, membrane stiffness has been used as an indicator of plasma membrane tension. SICM can be used to measure membrane stiffness without directly contacting the membrane and has been used to estimate membrane tension ^24,25^. In SICM, a glass nanopipette is used as a probe to monitor ion currents between an Ag/AgCl electrode placed inside the pipette and an Ag/AgCl electrode located in a bath. A constant pressure is applied to the pipette to generate a microfluidic flow, which deforms the cell surface. As the pipette approaches the cell, the ion current decreases. This decrease is slower for softer cells because the microfluidic flow progressively indents the cell surface. Therefore, the plasma membrane stiffness can be determined based on the ion current approach curve ^24^.

The plasma membrane tension was measured using a homemade SICM system with an inverted optical microscope (Eclipse-Ti, Nikon). The setup of the SICM system, the scanning algorithms and the protocol were described previously ^40^. Before the measurements were acquired, the culture medium was replaced with L-15 medium containing 2% B-27 supplement, 1 mM glutamine and 100 U/mL penicillin‒streptomycin. Glass nanopipettes (aperture inner radius 100 nm) filled with L-15 medium were used as probes. Scanning was performed in hopping mode with the following parameters: hopping amplitude, 3-5 μm; waiting time after lateral movement, 1 ms (during this time, a reference current was measured as the average of the direct current through the probe); probe approach and withdrawal speed, 20 and 500-1000 nm/ms, respectively; applied pressure, 10 kPa; and set point, 98% of the reference current. We recorded ion currents and the vertical probe position as the probe approach the cell surface. For the experiments shown in Figure 1D, we obtained data for a 25 x 25 µm^2^ area containing 128 x 128 points. For the experiments shown in Figure 1E and 1F, we obtained data for a 10 x 10 µm^2^ area containing 32 x 32 points, and repeated the experiment 3 times for each cell at 1 h intervals. After the measurements were performed, optical cell images were acquired using a CSI S Plan Fluor ELWD 40x/0.6 NA objective lens (Nikon), a complementary metal oxide semiconductor (CMOS) camera (ORCA Flash 4.0 C13440-20CU, Hamamatsu) and imaging software (HCI image, Hamamatsu) or using a Plan Flour 10x/0.30 NA objective lens (Nikon), a CMOS camera (Wraycam-noa2000, Wraymer) and imaging software (MicroStudio, Wraymer). The SICM data were analyzed and stiffness maps were produced using a handmade program developed based on LabVIEW2014 (National Instruments). The approach curve of each point was plotted as the ion current versus the vertical probe position, and the slope between 98.5% and 99% of the reference current was determined with a line fit. Young’s modulus of the cell surface was determined as described previously^24^:

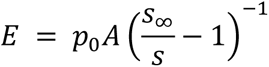

where *E* is Young’s modulus, *p*_0_ is the applied pressure, *A* is the pipette geometrical parameter (whose value is 0.4, as determined previously ^24^), and *s* and *s*_∞_ are the measured slopes on the cell surface and substrate, respectively.

### Ca^2+^ imaging

[Ca^2+^]_i_ was monitored using the ratiometric calcium indicator dye CalRed R525/650 AM (AAT Bioquest); when this indicator is excited at 488 nm upon Ca^2+^ binding, the emission signals increased at 525 nm and decreased at 650 nm. After 2 days in culture, olfactory interneurons were loaded with 1 µM CalRed R525/650 AM, 0.1% DMSO (Sigma) and 0.04% F-127 (AAT Bioquest) diluted in the culture medium for 1 h at 37°C and 5% CO_2_. The cells were subsequently washed with the culture medium. Before imaging, the culture medium was replaced with prewarmed artificial cerebrospinal fluid (ACSF) (10 mM HEPES, pH 7.4, 125 mM NaCl, 25 mM glucose, 2.5 mM KCl, 2 mM CaCl_2_, 1 mM MgCl_2_). Imaging was performed at 37°C using a confocal microscope (LSM710, Carl Zeiss) equipped with a Plan-Apochromat 20x/0.8 objective lens. The dye-loaded olfactory interneurons were excited with a 488 nm argon laser, and the emission signals were detected simultaneously at 525 ± 25 nm and > 611 nm. Differential interference contrast (DIC) and fluorescence images were acquired at 5-sec intervals for 15 min or 30 min. To image cells expressing miRNA, EBFP fluorescence was used as an indicator of transfected cells. [Ca^2+^]_i_ analyses were performed using ImageJ software (NIH). The fluorescence intensities at 525 nm and 650 nm in the same region were measured over time at the soma. [Ca^2+^]_i_ was calculated as the ratio of the fluorescence intensity at 525 nm to the fluorescence intensity at 650 nm. The average of [Ca^2+^]_i_ was set as the baseline (F0), and the [Ca^2+^]_i_ in each frame (F) was normalized to the baseline (F/F_0_). A change in [Ca^2+^]_i_ was judged as a Ca^2+^ transient if F/F_0 >_ 40%.

### Mechanical stretching of the leading process

Neurons suspended in a mixture of 75% Matrigel and 25% L-15 medium were plated on elastic chambers and cultured for 2 days as described above. For Ca^2+^ imaging, after a control image was acquired, the chamber was set on the stretching device (Strex, catalog number STB-CH-04ST-20) and stretched uniaxially (20% stretch) (Figure S1A). We then identified the same cell and acquired Ca^2+^ images after the stretching process. We also measured the angle of the leading process with respect to the stretch direction (*θ*) (Figure S1B). When the angle was between 0° and 30°, we determined that the leading process was extended (parallel extension). On the other hand, when the angle was between 60° and 90°, the stretch direction was determined to be perpendicular to the leading process. For immunocytochemical analysis, olfactory interneurons were subjected to 20% uniaxial stretch for 15 min at 37°C before fixation.

### Analyses of neuronal migration in Matrigel

Before observation, the culture medium was replaced with L-15 medium containing 2% B- 27 supplement, 1 mM glutamine and 100 U/mL penicillin‒streptomycin. Phase contrast images were acquired every 2 min for 2 h at 37°C using a fluorescence microscope (IX81, Olympus) equipped with Plan Fluor 40x, 0.60 NA (Olympus) and an electron multiplying- CCD camera (EMCCD, iXon DU888, Andor) and MetaMorph. To image the cells expressing miRNA, mCherry fluorescence was used as an indicator of transfected cells. Time-lapse images were analyzed and quantified using ImageJ and the manual tracking plugin. The migration speed (Figure 4E) was calculated by dividing the migration distance of the soma by the observation time. When the soma translocated any distance during the 2-min image acquisition intervals, we judged that neurons were in the somal translocation phase. The somal translocation speed (Figure 4D) was calculated by dividing the migration distance of the soma during the somal translocation phase by the observation time.

### qPCR analysis of migrating neurons

SVZ explants were prepared and cultured as previously described ^15^. Briefly, the SVZs were dissected from the P5 mouse brains and cut into blocks 100-150 μm in diameter. The SVZ tissue blocks were then embedded in a mixture of 75% Matrigel and 25% L-15 medium, and cultured in neurobasal medium containing 2% B-27 supplement, and 1 mM glutamine in a humidified 5% CO_2_ incubator at 37°C. After 2 days in culture, the SVZ blocks were removed from the olfactory interneurons that migrated out from the blocks. Then, the total RNA of the migrating neurons was isolated using an RNeasy mini kit (Qiagen) according to the manufacturer′s protocol followed by DNaseI digestion (Thermo Fisher Scientific, catalog number 11766051). The RNA was then reverse transcribed into cDNA using Superscript IV VILO master mix (Thermo Fisher Scientific) according to the manufacturer′s protocol, and cDNA was used in the qPCR assays. qPCR assays were performed using the LightCycler 96 system (Roche), KAPA SYBR FAST qPCR kit (Nippon Genetics) and the following primer sets: Piezo1, 5′- TTGACCCTGCCAACTGGTTT-3′ and 5′-GCCTCAAACACCAGCAACAG-3′; Piezo2, 5′-GTATTGGATCTACGTCTGCGG-3′ and 5′-CAGGATCTTCCTCCACCACTCG-3′; TRPC1, 5′-CAGAAGGACTGTGTGGGCAT-3′ and 5′- GAACAGAGCAAAGCAGGTGC-3′; TRPC3, 5′-AGTGTCTGGTCGTGTTGGTC-3′ and 5′-AAGAATCTTCCCCAGCCTGC-3′; TRPC5, 5′- TACCAATGTGAAGGCCCGAC-3′ and 5′-TATCAGCATGATCGGCAATG-3′; TRPC6, 5′-GCCGTCCAAATCTCAGCCGTT-3′ and 5′- TGGACAGGAGCTGTTGCTGAC-3′; TRPV2, 5′-GTTTGACCGTGACCGACTCT-3′ and 5′-CGTCTTTCCAGTGGAGCCTT-3′; TRPV4, 5′-GCTTCTTCCAGCCCAAGGAT-3′ and 5′-TGTCGCCTCATGTCAGCTTT-3′; TRPA1, 5′- ACAAGAAAGCCAGCCCTCTC-3′ and 5′-GGTTGCAGCAAAATGGAGGG-3′; TACAN, 5′-CTGCAGCAGGACTTCCAAGGT-3′ and 5′- CAGTTGGCCTGGAGTTTGGTC-3′; Tmem63a, 5′-CAGGGCAACAAGACTTTGAA- 3′ and 5′-TGTGCCTCTGAAAAGACAGG-3′; Tmem63b, 5′- AGAACAGGACGACCATGCACA-3′ and 5′-AGAGCCAGCGGAAGAAGAGGT-3′; GAPDH, 5′-TGATGGGTGTGAACCACGAG-3′ and 5′- GGCATGGACTGTGGTCATGA-3′.

### Preparation of cell lysate and immunoblotting

Cultured neurons were lysed with RIPA buffer (50 mM Tris-HCl (pH 8.0), 1 mM EDTA, 150 mM NaCl, 1% Triton X-100, 0.1% SDS, 0.1% sodium deoxycholate, 1 mM DTT, 1 mM PMSF, and 0.01 mM leupeptin), and incubated for 2 h at 4°C. The cell lysate was then centrifuged at 17900 x *g* for 10 min at 4°C. The supernatant was mixed with an equal volume of 2 x SDS sample buffer (131 mM Tris-HCl, pH 6.8, 21% glycerol, 4% SDS, 0.05% bromophenol blue and 5% β-mercaptoethanol). The mixture was incubated for 2 h at 37°C, followed by SDS-polyacrylamide gel electrophoresis. Immunoblotting was performed as previously described ^15^. The following primary antibodies were used in immunoblotting: rabbit anti-Tmem63b (1:200) (Novus Biologicals, catalog number NBP2- 58473), mouse anti-actin (1:10000) (Millipore, catalog number MAB1501R). The following secondary antibodies were used for immunoblotting: HRP-conjugated donkey anti-rabbit IgG (1:2000) (GE Healthcare, catalog number NA934) and HRP-conjugated donkey anti-mouse IgG (1:5000) (Bio-Rad, catalog number 1706516).

### Immunocytochemical analysis

Olfactory interneurons subjected to 20% uniaxial stretch were fixed with 4% paraformaldehyde dissolved in Krebs buffer (118 mM NaCl, 4.7 mM KCl, 1.2 mM KH_2_PO_4_, 1.2 mM MgSO_4_, 4.2 mM NaHCO_3_, 2 mM CaCl_2_, 10 mM glucose, 400 mM sucrose, and 10 mM HEPES (pH 7.0)) for 10 min on ice, followed by treatment with 0.1% Triton X-100 in phosphate-buffered saline (PBS) for 15 min on ice and 10% fetal bovine serum in PBS for 1 h at room temperature. They were then incubated with primary antibody diluted in PBS containing 10% fetal bovine serum overnight at 4°C. The following primary antibodies were used: rabbit anti-myosin light chain 2 (1:500) (Abcam, catalog number ab79935) and mouse anti-phospho-myosin light chain 2 (Ser19) (1:500) (Cell Signaling Technology, catalog number 3675). Cells were washed with PBS, and then incubated with secondary antibody and 1 μg/mL DAPI (Thermo Fisher Scientific) diluted in PBS for 1 h at room temperature. The following secondary antibodies were used: Alexa Fluor 647- conjugated donkey anti-rabbit IgG (1:1,000) (Abcam, catalog number ab150075) and Alexa Fluor 488 conjugated goat anti-mouse (1:1,000) (Thermo Fisher Scientific, catalog number A-11029). The immunostained cells were mounted with 50% (v/v) glycerol (Nacalai Tesque) in PBS. Fluorescence images were acquired using a confocal laser microscope (Stellaris 8, Leica) equipped with an HC Plan-Apochromat 20x, 0.75 NA objective lens (Leica) and imaging software (LAS X, Leica). To acquire images of the entire olfactory interneurons, we obtained 6-15 confocal images at 2 μm intervals in the Z axis direction. Then, the confocal images were volume-stacked, and the fluorescence signals were quantified using ImageJ.

### Chemicals

The following reagents were used at the indicated concentrations in this study: 5 µM GsMTx4 (Tocris Bioscience), 100 µM blebbistatin (Sigma) and 10 µM ML7 (Tocris Bioscience). A stock solution of GsMTx4 was prepared in H2O. Stock solutions of blebbistatin and ML7 were prepared in DMSO.

## QUANTIFICATION AND STATISTICAL ANALYSIS

Statistical analyses were performed using Excel 2016 (Microsoft) or GraphPad Prism 7 (GraphPad Software). For samples with more than 7 data points, the D′Agostino–Pearson normality test was used to determine whether the data followed a normal distribution. For cases in which the number of data points was between 3 and 7, the Shapiro‒Wilk test was used for the normality test. We also tested the equality of variation with the F test for two independent groups that followed normal distributions. Significance tests were performed as follows: (1) two-tailed paired *t*-test to compare normally distributed data from two dependent groups; (2) two-tailed Wilcoxon signed-rank test to compare nonnormally distributed data from two dependent groups; (3) two-tailed unpaired Student′s *t*-test to compare normally distributed data with equal variance from two independent groups; (4) two-tailed unpaired Welch′s *t*-test to compare normally distributed data with unequal variance from two independent groups; (5) two-tailed Mann–Whitney *U*-test to compare nonnormally distributed data from two independent groups; (6) two-tailed one-way ANOVA with Tukey′s post hoc test to compare normally distributed data from three groups; (7) two-tailed Kruskal‒Wallis tests with Dunn′s multiple comparison test to compare nonnormally distributed data from three groups; and (8) two-tailed Spearman’s correlation coefficient test to analyze nonnormally distributed data. The statistical information and number of samples for each experiment are indicated in the figure legends. For detailed statistical results including the test statistics and exact p values, see the statistical source data associated with each figure. All data are shown as the mean ± SEM. Statistical significance was defined as ***p < 0.01; **p < 0.02; *p < 0.05; ns, not significant. For the experiments using shootin1 KO mice, littermates were allocated into the experimental groups by genotyping. For the other experiments, sample allocation was randomized as cells were derived from the same resource. The numbers for the groups of each sample were based on those in previously published studies. No data were excluded if the experiments were successfully performed. All experiments were performed at least three times and reliably reproduced. Investigators were blind to the experimental groups for each analysis, except biochemical analyses.

## SUPPLEMENTAL VIDEO AND TABLE LEGENDS

**Video S1. Related to Figure 2. Time-lapse movies of DIC (left) and [Ca^2+^]_i_ (right) of a migrating olfactory interneuron loaded with 1 µM CalRed R525/650 (see Figure 2A and B).** Images were acquired at 5-sec intervals for 30 min. Scale bar, 10 µm.

**Video S2. Related to Figure 2. A time-lapse movie of [Ca^2+^]_i_ (fire) of migrating olfactory interneuron loaded with 1 µM CalRed R525/650 (see Figure 2H).** The neuron was cultured on an elastic chamber and exposed to 20% stretch (yellow arrows). Images were acquired at 5-sec intervals for 15 min before and after 20% stretch. Scale bar, 10 µm.

**Video S3. Related to Figure 4. Time-lapse movies of migrating olfactory interneuron expressing control miRNAi (left), Tmem63b miRNAi (middle) and Tmem63b miRNA + Tmem63b^r^ (right) (see Figures 4A and B).** Phase contrast images were acquired at 2- min intervals for 120 min. Scale bar, 10 µm.

**Table S1. Related to Figure 3. qPCR primer sets (see Figure 3B).**

## REFERENCES

1. Ross, M.E., and Walsh, C.A. (2001). Human brain malformations and their lessons for neuronal migration. Annu Rev Neurosci 24, 1041–1070. 10.1146/annurev.neuro.24.1.1041.

2. Hatten, M.E. (2002). New directions in neuronal migration. Science 297, 1660–1663. 10.1126/science.1074572.

3. Rakic, P. (2009). Evolution of the neocortex: a perspective from developmental biology. Nat Rev Neurosci 10, 724–735 10.1038/nrn2719.

4. Saito, K., Okamoto, M., Watanabe, Y., Noguchi, N., Nagasaka, A., Nishina, Y., Shinoda, T., Sakakibara, A., and Miyata, T. (2019). Dorsal-to-ventral cortical expansion is physically primed by ventral streaming of early embryonic preplate neurons. Cell Rep 29, 1555–1567.e1555. 10.1016/j.celrep.2019.09.075.

5. Nakajima, C., Sawada, M., and Sawamoto, K. (2021). Postnatal neuronal migration in health and disease. Curr Opin Neurobiol 66, 1–9. 10.1016/j.conb.2020.06.001.

6. Edmondson, J.C., and Hatten, M.E. (1987). Glial-guided granule neuron migration in vitro: a high-resolution time-lapse video microscopic study. J Neurosci 7, 1928–1934. 10.1523/jneurosci.07-06-01928.1987.

7. O’Rourke, N.A., Dailey, M.E., Smith, S.J., and McConnell, S.K. (1992). Diverse migratory pathways tn the developing cerebral cortex. Science 258, 299–302. doi:10.1126/science.1411527.

8. Marin, O., Valiente, M., Ge, X., and Tsai, L.H. (2010). Guiding neuronal cell migrations. Cold Spring Harb Perspect Biol 2, a001834. 10.1101/cshperspect.a001834.

9. Shu, T., Ayala, R., Nguyen, M.-D., Xie, Z., Gleeson, J.G., and Tsai, L.-H. (2004). Ndel1 operates in a common pathway with LIS1 and cytoplasmic dynein to regulate cortical neuronal positioning. Neuron 44, 263–277. 10.1016/j.neuron.2004.09.030.

10. Schaar, B.T., and McConnell, S.K. (2005). Cytoskeletal coordination during neuronal migration. Proc Natl Acad Sci U S A 102, 13652–13657. 10.1073/pnas.0506008102.

11. Solecki, D.J., Trivedi, N., Govek, E.E., Kerekes, R.A., Gleason, S.S., and Hatten, M.E. (2009). Myosin II motors and F-actin dynamics drive the coordinated movement of the centrosome and soma during CNS glial-guided neuronal migration. Neuron 63, 63–80. 10.1016/j.neuron.2009.05.028.

12. Zhang, X., Lei, K., Yuan, X., Wu, X., Zhuang, Y., Xu, T., Xu, R., and Han, M. (2009). SUN1/2 and Syne/Nesprin-1/2 complexes connect centrosome to the nucleus during neurogenesis and neuronal migration in mice. Neuron 64, 173–187. 10.1016/j.neuron.2009.08.018.

13. Martini, F.J., and Valdeolmillos, M. (2010). Actomyosin contraction at the cell rear drives nuclear translocation in migrating cortical interneurons. J Neurosci 30, 8660–8670. 10.1523/JNEUROSCI.1962-10.2010.

14. He, M., Zhang, Z.H., Guan, C.B., Xia, D., and Yuan, X.B. (2010). Leading tip drives soma translocation via forward F-actin flow during neuronal migration. J Neurosci 30, 10885–10898. 10.1523/JNEUROSCI.0240-10.2010.

15. Minegishi, T., Uesugi, Y., Kaneko, N., Yoshida, W., Sawamoto, K., and Inagaki, N. (2018). Shootin1b mediates a mechanical clutch to produce force for neuronal migration. Cell Rep 25, 624–639 e626. 10.1016/j.celrep.2018.09.068.

16. Komuro, H., and Rakic, P. (1996). Intracellular Ca2+ fluctuations modulate the rate of neuronal migration. Neuron 17, 275–285. 10.1016/s0896-6273(00)80159-2.

17. Wichterle, H., Garcia-Verdugo, J.M., and Alvarez-Buylla, A. (1997). Direct evidence for homotypic, glia-independent neuronal migration. Neuron 18, 779–791 10.1016/s0896-6273(00)80317-7.

18. Murthy, S.E., Dubin, A.E., Whitwam, T., Jojoa-Cruz, S., Cahalan, S.M., Mousavi, S.A.R., Ward, A.B., and Patapoutian, A. (2018). OSCA/TMEM63 are an evolutionarily conserved family of mechanically activated ion channels. Elife 7. 10.7554/eLife.41844.

19. Lamoureux, P., Buxbaum, R.E., and Heidemann, S.R. (1989). Direct evidence that growth cones pull. Nature 340, 159–162 10.1038/340159a0.

20. Houk, A.R., Jilkine, A., Mejean, C.O., Boltyanskiy, R., Dufresne, E.R., Angenent, S.B., Altschuler, S.J., Wu, L.F., and Weiner, O.D. (2012). Membrane tension maintains cell polarity by confining signals to the leading edge during neutrophil migration. Cell 148, 175–188. 10.1016/j.cell.2011.10.050.

21. Tsujita, K., Takenawa, T., and Itoh, T. (2015). Feedback regulation between plasma membrane tension and membrane-bending proteins organizes cell polarity during leading edge formation. Nat Cell Biol 17, 749–758 10.1038/ncb3162.

22. De Belly, H., Yan, S., Borja da Rocha, H., Ichbiah, S., Town, J.P., Zager, P.J., Estrada, D.C., Meyer, K., Turlier, H., Bustamante, C., and Weiner, O.D. (2023). Cell protrusions and contractions generate long-range membrane tension propagation. Cell 186, 3049–3061.e3015. 10.1016/j.cell.2023.05.014.

23. Colom, A., Derivery, E., Soleimanpour, S., Tomba, C., Molin, M.D., Sakai, N., Gonzalez- Gaitan, M., Matile, S., and Roux, A. (2018). A fluorescent membrane tension probe. Nat Chem 10, 1118–1125. 10.1038/s41557-018-0127-3.

24. Rheinlaender, J., and Schaffer, T.E. (2013). Mapping the mechanical stiffness of live cells with the scanning ion conductance microscope. Soft Matter 9, 3230–3236. 10.1039/c2sm27412d.

25. Bednarska, J., Pelchen-Matthews, A., Novak, P., Burden, J.J., Summers, P.A., Kuimova, M.K., Korchev, Y., Marsh, M., and Shevchuk, A. (2020). Rapid formation of human immunodeficiency virus-like particles. Proc Natl Acad Sci U S A 117, 21637–21646. 10.1073/pnas.2008156117.

26. Lee, J., Ishihara, A., Oxford, G., Johnson, B., and Jacobson, K. (1999). Regulation of cell movement is mediated by stretch-activated calcium channels. Nature 400, 382–386. 10.1038/22578.

27. Douguet, D., and Honore, E. (2019). Mammalian mechanoelectrical transduction: structure and function of force-gated ion channels. Cell 179, 340–354. 10.1016/j.cell.2019.08.049.

28. Jiang, J., Zhang, Z.H., Yuan, X.B., and Poo, M.M. (2015). Spatiotemporal dynamics of traction forces show three contraction centers in migratory neurons. J Cell Biol 209, 759–774. 10.1083/jcb.201410068.

29. Umeshima, H., Nomura, K.I., Yoshikawa, S., Horning, M., Tanaka, M., Sakuma, S., Arai, F., Kaneko, M., and Kengaku, M. (2019). Local traction force in the proximal leading process triggers nuclear translocation during neuronal migration. Neurosci Res 142, 38–48. 10.1016/j.neures.2018.04.001.

30. Tsujita, K., Satow, R., Asada, S., Nakamura, Y., Arnes, L., Sako, K., Fujita, Y., Fukami, K., and Itoh, T. (2021). Homeostatic membrane tension constrains cancer cell dissemination by counteracting BAR protein assembly. Nat Commun 12, 5930. 10.1038/s41467-021-26156- 4.

31. Gnanasambandam, R., Ghatak, C., Yasmann, A., Nishizawa, K., Sachs, F., Ladokhin, A.S., Sukharev, S.I., and Suchyna, T.M. (2017). GsMTx4: mechanism of inhibiting mechanosensitive ion channels. Biophys J 112, 31–45. 10.1016/j.bpj.2016.11.013.

32. Horigane, S.I., Ozawa, Y., Yamada, H., and Takemoto-Kimura, S. (2019). Calcium signalling: a key regulator of neuronal migration. J Biochem 165, 401–409. 10.1093/jb/mvz012.

33. Ofer, N., Mogilner, A., and Keren, K. (2011). Actin disassembly clock determines shape and speed of lamellipodial fragments. Proc Natl Acad Sci U S A 108, 20394–20399. doi:10.1073/pnas.1105333108.

34. Tsai, T.Y., Collins, S.R., Chan, C.K., Hadjitheodorou, A., Lam, P.Y., Lou, S.S., Yang, H.W., Jorgensen, J., Ellett, F., Irimia, D., et al. (2019). Efficient front-rear coupling in neutrophil chemotaxis by dynamic myosin II localization. Dev Cell 49, 189–205.e186. 10.1016/j.devcel.2019.03.025.

35. Valiente, M., and Marin, O. (2010). Neuronal migration mechanisms in development and disease. Curr Opin Neurobiol 20, 68–78. 10.1016/j.conb.2009.12.003.

36. Evsyukova, I., Plestant, C., and Anton, E.S. (2013). Integrative mechanisms of oriented neuronal migration in the developing brain. Annu Rev Cell Dev Biol 29, 299–353. 10.1146/annurev-cellbio-101512-122400.

37. Stouffer, M.A., Golden, J.A., and Francis, F. (2016). Neuronal migration disorders: Focus on the cytoskeleton and epilepsy. Neurobiol Dis 92, 18–45. 10.1016/j.nbd.2015.08.003.

38. Baba, K., Yoshida, W., Toriyama, M., Shimada, T., Manning, C.F., Saito, M., Kohno, K., Trimmer, J.S., Watanabe, R., and Inagaki, N. (2018). Gradient-reading and mechano- effector machinery for netrin-1-induced axon guidance. Elife 7, e34593. 10.7554/eLife.34593.

39. Diz-Munoz, A., Fletcher, D.A., and Weiner, O.D. (2013). Use the force: membrane tension as an organizer of cell shape and motility. Trends Cell Biol 23, 47–53. 10.1016/j.tcb.2012.09.006.

40. Takahashi, Y., Zhou, Y., Miyamoto, T., Higashi, H., Nakamichi, N., Takeda, Y., Kato, Y., Korchev, Y., and Fukuma, T. (2020). High-speed SICM for the visualization of nanoscale dynamic structural changes in hippocampal neurons. Anal Chem 92, 2159–2167. 10.1021/acs.analchem.9b04775.

