## Supplemental Information for "Mechanical signaling through membrane tension induces somal translocation during neuronal migration"

**This PDF file includes:**

Figures S1-S4

**Other supplemental information for this manuscript include:**

Table S1

Supplemental Videos 1-3

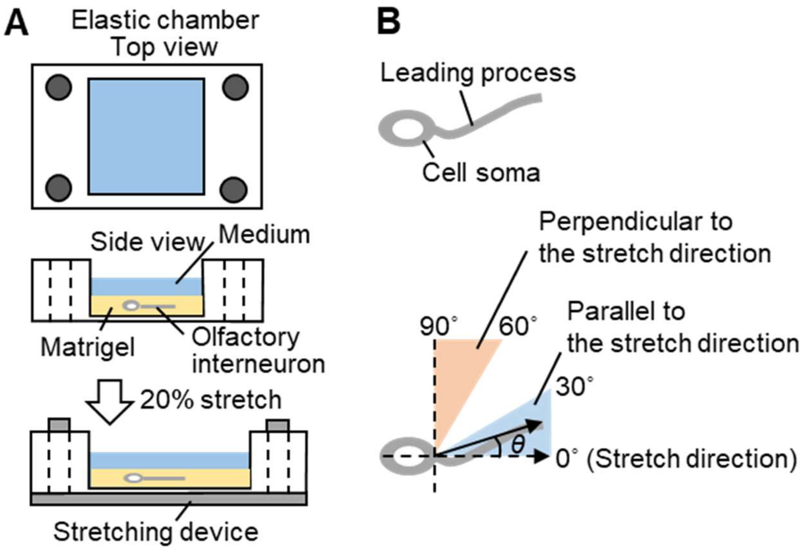


**Figure S1. Related to Figure 2. Schematic diagrams of cell stretching assay.**

(A) Olfactory interneurons were suspended in Matrigel, and plated on elastic chambers. The chamber was set on the stretching device, and stretched uniaxially (20% stretch).

(B) The definition of cell orientation in this study. We measured the angle of the leading process with respect to the stretch direction ($\theta$). When the angle was between 0° and 30°, we determined that the leading process was extended (parallel extension). On the other hand, when the angle was between 60° and 90°, the stretch direction was determined to be perpendicular to the leading process.

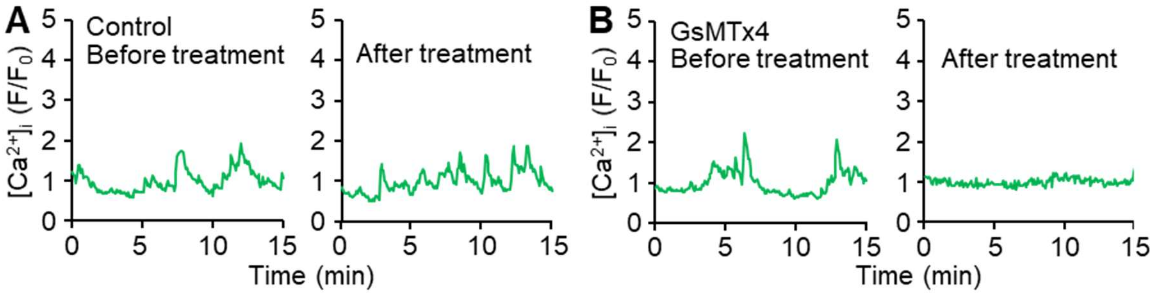


**Figure S2. Related to Figure 3. Inhibition of mechanosensitive channels reduces Ca2+ transients of migrating olfactory interneurons.**

(A and B) Time courses of $[Ca^{2+}]_i$ in migrating olfactory interneurons treated with vehicle control (A) or 5 μM GsMTx4 (B). Migrating neurons were imaged at 5-sec intervals for 15 min before and after treatment.

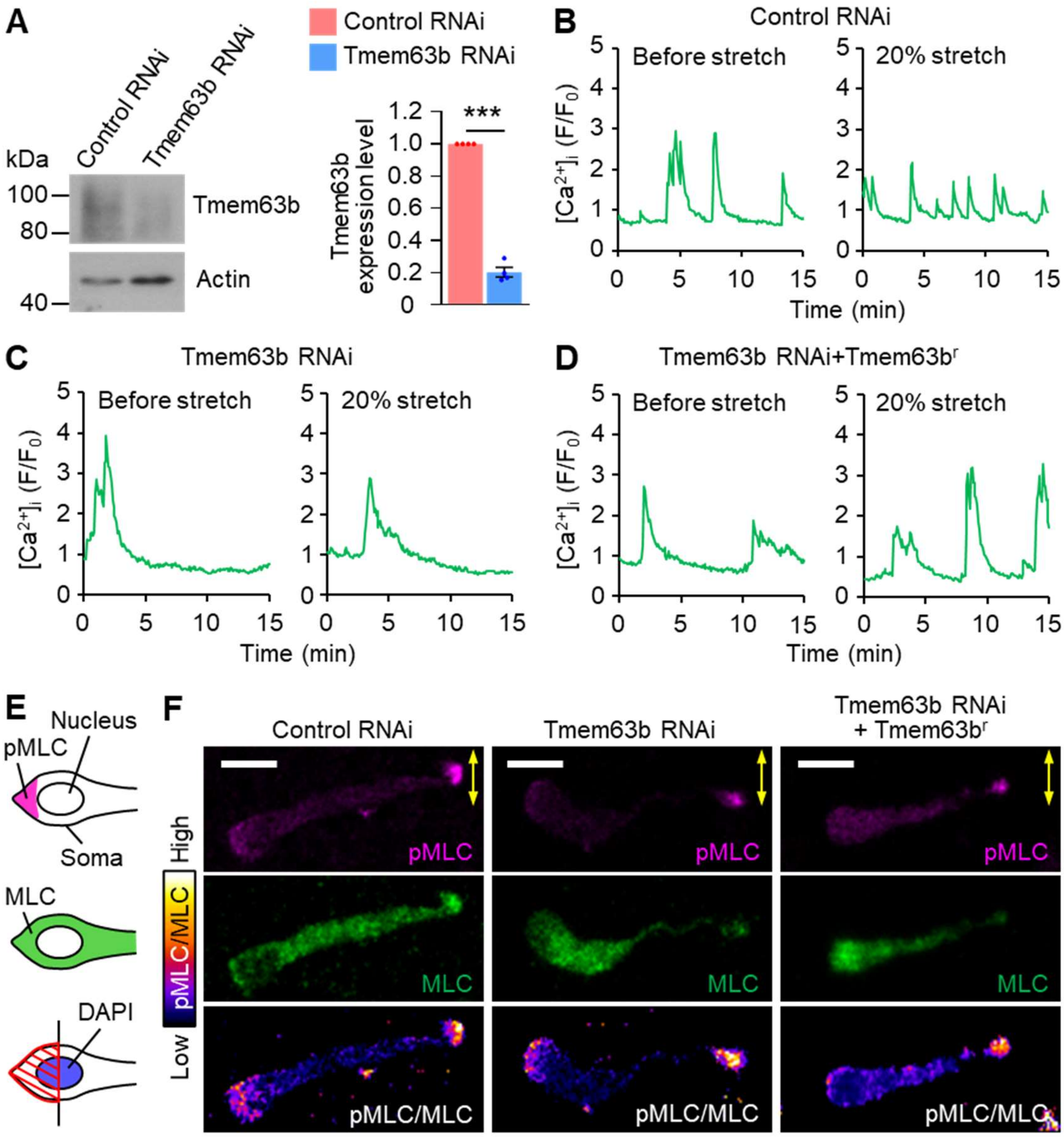


**Figure S3. Related to Figure 3. Analyses of extension-induced $Ca^{2+}$ transients and myosin II activity in olfactory interneurons expressing control RNAi, Tmem63b RNAi or Tmem63b RNAi + Tmem63b$^r$.**

(A) Immunoblot analysis of Tmem63b in olfactory interneurons expressing control miRNA or Tmem63b miRNA. Anti-actin antibody served as loading controls. The graph shows the Tmem63b expression levels. The level of Tmem63b in olfactory interneurons expressing control miRNA was normalized as 1. n = 4 independent experiments. Data represent means ± SEM. Statistical analysis was performed using the two-tailed unpaired Welch's t-test. ***p < 0.01.

(B-D) Time courses of $[Ca^{2+}]_i$ in migrating olfactory interneurons expressing control miRNA (B), Tmem63b miRNA (C) or Tmem63b miRNA + RNAi-refractory Tmem63b (Tmem63b$^r$) (D). Migrating olfactory interneurons cultured on elastic chambers were imaged at 5-sec intervals for 15 min before (control) and after uniaxial stretch (20% stretch). We imaged olfactory interneurons with leading processes oriented parallel to the stretch direction. Immunocytochemical analysis of myosin II activity.

(E) A diagram describing the analysis of myosin II activity in olfactory interneurons. Fluorescent images of pMLC (magenta), MLC (green) and DAPI (blue) in olfactory interneurons were acquired. We then drew a line that cross the middle of nucleus stained by DAPI, and measured the fluorescent intensities of pMLC and MLC within the posterior half of the soma (shaded area). The myosin II activity was calculated as the ratio of the fluorescence intensity of pMLC to that of MLC.

(F) Fluorescent images of olfactory interneurons expressing control miRNA, Tmem63b miRNA or Tmem63b miRNA + Tmem63br. Olfactory interneurons cultured on elastic chambers were fixed after 20% stretch, and then stained with anti-phospho-myosin light chain 2 (pMLC) antibody (magenta), anti-myosin light chain 2 (MLC) antibody (green) and DAPI. The ratio of the fluorescence intensity of pMLC to that of MLC (pMLC/MLC) was displayed by the pseudocolor bar. Neurons with leading processes oriented perpendicular to the stretch direction (yellow arrows) were analyzed as controls of the data in Figure 3D. Scale bars, 10 μm.

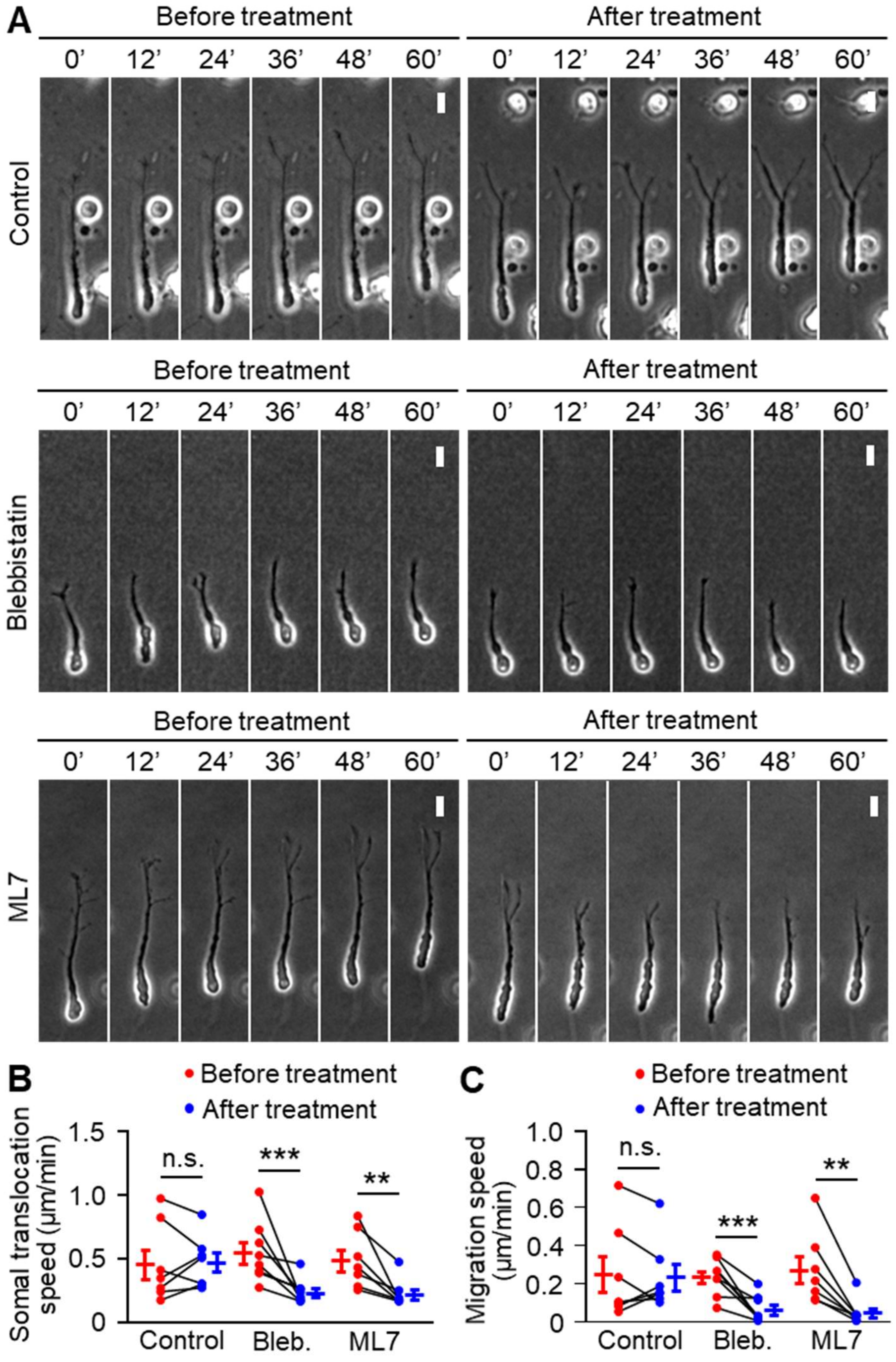


**Figure S4. Related to Figure 4. Inhibition of myosin II or myosin light chain kinase decreases somal translocation and migration speeds of olfactory interneurons.**

(A) Time-lapse images of migrating olfactory interneurons before and after treatment with DMSO (control), 100 μM blebbistatin or 10 μM ML7. Neurons were imaged at 2-min intervals.

(B and C), Somal translocation speed (B) and migration speed (C) of migrating olfactory interneurons treated with DMSO (control), 100 μM blebbistatin (bleb.) or 10 μM ML7 in (A). Control, $n = 7$ cells; blebbistatin, $n = 8$ cells; ML7, $n = 7$ cells. Data represent means ± SEM. Two-tailed Wilcoxon signed-rank test (B and C). **$p < 0.02$; ***$p < 0.01$; n.s., not significant. Scale bars, 10 μm.
